# *Pseudomonas aeruginosa* leucine aminopeptidase influences early biofilm composition and structure via vesicle-associated anti-biofilm activity

**DOI:** 10.1101/784918

**Authors:** Caitlin N. Esoda, Meta J. Kuehn

## Abstract

*Pseudomonas aeruginosa*, known as one of the leading causes of disease in cystic fibrosis (CF) patients, secretes a variety of proteases. These enzymes contribute significantly to *P. aeruginosa* pathogenesis and biofilm formation in the chronic colonization of CF patient lungs, as well as playing a role in infections of the cornea, burn wounds and chronic wounds. We previously characterized a secreted *P. aeruginosa* peptidase, PaAP, that is highly expressed in chronic CF isolates. This leucine aminopeptidase is highly expressed during infection and in biofilms, and it associates with bacterial outer membrane vesicles (OMVs), structures known to contribute to virulence mechanisms in a variety of Gram-negative species and one of the major components of the biofilm matrix. We hypothesized that PaAP may play a role in *P. aeruginosa* biofilm formation. Using a lung epithelial cell/bacterial biofilm coculture model, we show that PaAP deletion in a clinical *P. aeruginosa* background alters biofilm microcolony composition to increase cellular density, while decreasing matrix polysaccharide content, and that OMVs from PaAP expressing strains but not PaAP alone or in combination with PaAP deletion strain-derived OMVs could complement this phenotype. We additionally found that OMVs from PaAP expressing strains could cause protease-mediated biofilm detachment, leading to changes in matrix and colony composition. Finally, we showed that the OMVs could also mediate the detachment of biofilms formed by both non-self *P. aeruginosa* strains and *Klebsiella pneumoniae*, another respiratory pathogen. Our findings represent novel roles for OMVs and the aminopeptidase in the modulation of *P. aeruginosa* biofilm architecture.

**Importance:** Biofilm formation by the bacterial pathogen *P. aeruginosa* is known to contribute to drug-resistance in nosocomial infections and chronic lung infections of cystic fibrosis patients. In order to treat these infections more successfully, the mechanisms of bacterial biofilm development must be elucidated. While both bacterially-secreted aminopeptidase and outer membrane vesicles have been shown to be abundant in *P. aeruginosa* biofilm matrices, the contributions of each of these factors to the steps in biofilm generation have not been well studied. This work provides new insight as to how these bacterial components mediate the formation of a robust, drug-resistant extracellular matrix and implicates outer membrane vesicles as active components of biofilm architecture, expanding our overall understanding of *P. aeruginosa* biofilm biology.

## Introduction

The Gram-negative bacterium *Pseudomonas aeruginosa* is a prominent opportunistic pathogen capable of causing both acute and chronic disease in a variety of compromised hosts. *P. aeruginosa* is most well known as a leading cause of morbidity and mortality in cystic fibrosis (CF) patients^1^. In these individuals, the pathogen establishes chronic, biofilm-based infections that may persist for decades in the unique, mucous-rich environment of the CF lung. In addition to lung tissue, *P. aeruginosa* forms biofilms on a wide range of substrates relevant to human infection, including corneal and skin tissue, as well as soil and water reservoirs and hospital surfaces, which can contribute to infection initiation and spread^2^.

*P. aeruginosa* is considered a model organism for the study of biofilm formation, and many of the cellular and matrix components contributing to this mode of growth have been studied previously^3–5^. The process of forming biofilm communities has also been characterized, revealing distinct growth phases. First the bacteria settle and attach onto a suitable host tissue or abiotic surface, then they begin to form stationary microcolonies and secrete extrapolymeric substance (EPS) to form a dense matrix. This matrix contains a variety of polysaccharides, with Psl and Pel polysaccharides present in all strains, and alginate found in mucoid *P. aeruginosa* isolates^6^. It also contains proteins, lipids, extracellular outer membrane vesicles (OMVs), and extracellular DNA (eDNA)^4, 5, 7^; however, the mechanisms by which many of these secreted bacterial components specifically affect microcolony growth and development remain unclear.

As with other extracellular pathogens, *P. aeruginosa* secretes many of its virulence determinants into the extracellular milieu to affect interactions with the host, including those in the biofilm matrix. Interestingly, many of the virulence factors secreted specifically by *P. aeruginosa* demonstrate proteolytic activities^8–10^. These enzymes, including elastase, protease IV, and alkaline protease, act on both bacterial and host proteins to directly impact host-pathogen interactions. For example, purified elastase has been found to cleave fibrin, laminin, various immunoglobulins, and components of the complement system^11^. Their activity not only interferes with host defense mechanisms but may also compromise host epithelial junctions, allowing the bacteria entrance to otherwise inaccessible tissues. These proteases can additionally facilitate biofilm formation and antibiotic resistance^12^. Based on the importance of known virulence-associated, secreted proteases in *P. aeruginosa* biofilm formation, it follows that other proteases in the secretome, many of which have not been fully characterized, may also play a role in these processes.

With the goal of identifying factors that may contribute to chronic infections, our lab compared secretome protein expression profiles of clinical *P. aeruginosa* isolates from chronically infected CF patients to those of highly passaged laboratory and environmental strains. Several proteins were identified as being more highly expressed in all the clinical strains, indicating their potential involvement in virulence and chronic infection mechanisms. Among the proteins identified was aminopeptidase PA2939, known as PaAP (***<u>P</u>****seudomonas **<u>a</u>**eruginosa* **<u>a</u>**mino**<u>p</u>**eptidase)^13^.

Cahan et al. first described the proteolytic activity of PaAP in *P. aeruginosa* supernatants and classified it as a leucine aminopeptidase and a member of the M28 family of metalloproteases^10^. PaAP expression is regulated through quorum sensing (QS) mechanisms, and the protein is secreted via the *Pseudomonas* type II Xcp secretion pathway^14^. PaAP enters the extracellular environment as a 536 amino acid proenzyme. Its C terminus is then cleaved by a variety of extracellular proteases to activate its enzymatic activity, and subsequently its N-terminus is auto-processed to create a 56 kDa, enzymatically active product^15^. It can be further processed extracellularly into several smaller, active products ranging from 55 kDa to 26 kDa. Outside of the cell, PaAP is found both soluble in the extracellular environment and as an abundant component of outer membrane vesicles (OMVs)^13^.

OMVs are formed from the envelope of Gram-negative bacteria and are small, discrete extracellular structures containing distinct membrane protein, lipid, and soluble periplasmic content^16^. They are known to interact with the external environment, including host tissues^17^. Vesicles produced by many Gram-negative pathogens, including *P. aeruginosa*, can elicit host immune responses, and many also serve as delivery mechanisms for bacterial virulence factors^17^. Of particular importance to our studies, OMVs have been implicated in the process of biofilm formation^7, 16, 18^. Schooling *et al.* demonstrated that OMVs are present in *P. aeruginosa* biofilm matrix and that bacteria in biofilms exhibit higher OMV production rates as compared to planktonic cells^19^. When these biofilm-derived vesicles were analyzed, PaAP was found to be one of the most abundant proteins^20^.

Bacterial density, as communicated through quorum sensing, is known to be an important regulatory facet of biofilm development^21^. Not only is PaAP expression regulated by quorum sensing, the XCP type II secretion system that secretes PaAP is also QS-regulated^14^. In addition, when *P. aeruginosa* was cultured on host epithelial cells, PaAP expression was found to increase significantly^22^ suggesting that PaAP may play a role in biofilm formation on the host cells. Together, these data suggest PaAP’s potential role in fine-tuned pathogenesis processes, including biofilm formation and infection.

In this study, we report that the formation and composition of early biofilms are modulated by novel roles for the secreted *P. aeruginosa* leucine aminopeptidase and secreted vesicles. Using a coculture model in which single-species bacterial biofilms are formed on a confluent layer of host cells, we show that PaAP expression inhibits the formation of cellularly dense microcolonies while promoting greater exopolysaccharide production and colistin resistance. Additionally, we show a secreted biofilm detachment activity that is mediated by *P. aeruginosa* vesicles and depends on vesicle-associated protease activity. These data provide novel insight into how secreted bacterial products modulate clinically-relevant bacterial biofilms.

## Materials and Methods

### Bacterial strains and cell culturhe methods

*P. aeruginosa* strains used include the laboratory strain PAO1 (Pf1 phage-cured from our lab collection), PA14 and associated mutants from a previously described transposon insertion library^23^, and minimally passaged, non-mucoid cystic fibrosis clinical isolate S470 along with its previously-described isogenic ΔPaAP (PA2939 deletion) mutant^13, 24^. To generate fluorescent strains, we transformed bacteria with the dTomato expression plasmid, p67T1^25^. *Klebsiella pneumoniae* strain 43816 was obtained from ATCC. A549 human lung epithelia carcinoma cells (ATCC CCL-185) were grown in F-12K media containing 10% fetal bovine serum plus penicillin/streptomycin/fungizone. Unless specifically mentioned, all reagents were obtained from Sigma-Aldrich.

### Coculture assay

A549 cells were grown in MatTek glass bottom 3.5 mm tissue culture treated dishes for 10 days at 37°C in 5% CO_2_, with media changes every two days. Bacterial cultures were grown overnight in LB (lysogeny broth) with the appropriate antibiotics, 1 mL of the culture was pelleted and resuspended in 1 mL sterile PBS (137 mM NaCl, 2.7 mM KCl, 10 mM Na_2_HPO_4_, 2 mM KH_2_PO_4_, pH 7.4). To fully resuspend the bacterial samples and reduce aggregation, each was passed through a 1 inch, 26-1/2G needle tip, 10 times. The OD_600_ was measured and conversion values were calculated to determine the number of cells/OD_600_ for each strain. For all following experiments, the OD_600_ was used to determine cell counts and multiplicity of infection (MOI). The bacterial samples were diluted in F-12K media (without antibiotic/antimycotic) and added to the A549-containing wells at MOI 30. The cocultures were incubated for 1h (37°C, 5% CO_2_) to allow the bacteria to attach to the cell surface. The cocultures were then washed three times with sterile PBS, and the media was replaced with F-12K containing 0.4% L-arginine. The cultures were incubated for an additional 4 h at 37°C and the media was replaced with microscopy-grade media (Sigma DMEM without Phenol Red, supplemented with 0.4% arginine). LDH release was measured using the CyQUANT LDH Cytotoxicity Assay kit (Thermo Fisher). Images were taken on a Zeiss 780 inverted confocal microscope. To ensure unbiased image collection, imaging fields were separated into evenly spaced sections and images were taken from a randomly selected spot within each section. To prevent photobleaching, no section was imaged more than once. Biomass calculations were completed using Comstat 2.1, and 3D colony models were created using the 3D viewer plugin for Image J. This protocol was adapted from previously published work^26^.

### Immunoblotting

To quantify PaAP protein in coculture biofilms, the cocultures were lysed with 100 µL RIPA buffer (VWR) and homogenized by repeated pipetting. For detection of PaAP in CFS by immunoblotting, 1 mL of CFS was used. All samples were precipitated with 250 µL trichloroacetic acid (TCA), incubated on ice for 1 h, pelleted by centrifugation (12,500 x g), washed with 1 mL acetone, re-pelleted (12,500 x g) and resuspended in 0.75 M Tris-Cl, pH 8.8, and 1x SDS-PAGE sample buffer (1% β-mercaptoethanol, 0.004% bromophenol blue, 6% glycerol, 2% sodium dodecyl sulfate (SDS), 50 mM Tris-Cl, pH 6.8), prior to SDS-PAGE on BioRad 4-20% Tris-HCl gels. For anti-PaAP immunoblotting, gels were transferred to Amersham Hybond 0.45 µm polyvinyldifluoride (PVDF) membranes (GE Healthcare), blocked with 1% non-fat dry milk in TBS (50 mM Tris-Cl, pH 7.6; 150 mM NaCl), and then incubated (4°C, overnight) with anti-PaAP antibody^15^ diluted 1:1000 in TBS with 0.1% Tween (TBST). Blots were washed three times with TBST, incubated (room temperature, 1 h) with goat anti-rabbit Li-Cor secondary antibody (Odyssey) diluted 1:40,000 TBST + milk, washed, and analyzed using an Li-Cor Odyssey CLx imaging system with Li-Cor Image Studio software. ImageJ was used to quantify the anti-PaAP reactive bands.

### qRT-PCR

Pellicle and coculture biofilms were grown for the indicated times as described. For coculture biofilms, supernatants were removed and 1 mL of Trizol was added to each sample to lyse both bacterial and A549 cells. For pellicle biofilms, 3 mL/well of cultures incubated in 6-well polystyrene dishes (VWR) were used to obtain sufficient biofilm material for analysis over the given time course. All samples were pipetted extensively to partially homogenize them and transferred to 15 mL conical tubes, in which the bacteria were pelleted. For planktonic culture comparisons, bacterial cultures were prepared as described for the coculture assay and grown in LB at 37°C with shaking for 5 h. Trizol (1 mL, Invitrogen) was used to lyse each sample. A standard Trizol extraction was performed as described by the Invitrogen protocol, and RNA was extracted from the aqueous phase. The RNA was reprecipitated using 3 M sodium acetate for further purification and resuspended in dH_2_0. RNA samples were then DNase treated and reverse transcribed using the Applied Biosystems High-Capacity cDNA Reverse Transcription kit protocol. Applied Biosystems Power Sybr Green PCR Master Mix was used to prepare the samples, and they were analyzed on a StepOne Plus real-time PCR machine (Applied Biosystems). ΔC_T_ values were calculated relative to *proC* as a housekeeping control, which was confirmed to have similar expression patterns in both biofilm and planktonic cultures at the concentrations of RNA used. Relative expression was calculated against planktonic S470 WT samples. Forward (F) and Reverse (R) primers used in this study are listed in **Table S1**.

### Antibiotic resistance

To determine antibiotic resistance of coculture biofilms and planktonic cultures, samples were grown as described above, and treated with the indicated concentrations of colistin hydrochloride (Sigma-Aldrich) for 2 h at 37°C. For cocultures, viability was determined by calculating fluorescent biomass retained at the cellular substrate, based on our finding that dead cells lost fluorescent activity. For planktonic cultures, viability was determined using the Live/Dead BacLight Bacterial Viability Kit (ThermoFisher Scientific). For detachment experiments, cocultures were treated with vesicles as described below, and supernatants were collected. The detached cells were pelleted, resuspended in PBS, incubated with the indicated colistin concentrations for 2 h at 37°C, and viability determined using BacLight staining.

### Pellicle biofilm quantitation

Bacterial cultures were grown overnight in LB (37°C, 200 rpm) with antibiotics if appropriate. The cultures were vortexed for 15 sec and diluted 1:1000 in Jensen’s media^27^, M63 minimal media (supplemented with 1 mM magnesium sulfate, 0.2% glucose, and 0.5% casamino acids as outlined previously^28^), LB+ 0.4% glucose, or tissue culture media (described above) with no antibiotics. Cultures were grown to an OD_600_ of 0.5 and diluted to OD_600_ of 0.1. Aliquots of the diluted culture (100 µL/well) were incubated in five 96-well flat bottom polystyrene dishes (VWR) for 0, 2, 6, 10, or 24 h at 37°C with no shaking and evaluated using a static biofilm quantitation assay^28^. Briefly, at the incubation endpoint, the plate was washed out using distilled water (dH_2_0). After three washes, 150 µL of 1% aqueous Crystal Violet solution was added to each well and the samples were incubated for 10 min at room temperature. The stained wells were washed three times using dH_2_0, 200 µL of 30% aqueous acetic acid was added to solubilize the dye, and the OD_495_ was measured.

### Pellicle biofilm imaging

Cultures were prepared as described above and diluted cultures were grown in 35 mm glass bottom dishes for 5 h at 30°C. Media was removed and the pellicles were stained using BacLight. The pellicles were imaged using a Zeiss 780 inverted confocal microscope and biomass was calculated using Comstat 2.1.

### Psl and matrix protein staining

To stain the biofilm extrapolymeric substance, 100 µg/mL HHA-FITC lectin (USA Biologicals) was added to cocultures 2 h before the imaging timepoint, and the samples were incubated at 37°C with 5% CO_2_. Pellicle samples were treated with 100 µg/mL HHA-TRITC (USA Biologicals) lectin for 2 h before the final imaging timepoint. For protein staining, pellicles were stained with Ruby FilmTracer SYPRO Ruby Biofilm Matrix Stain (ThermoFisher Scientific) for 5 min before imaging, as described in the reagent protocol. All samples were imaged using a Zeiss 780 inverted confocal microscope and staining was quantified using Comstat 2.1.

### OMV isolation and supernatant fractionation

Bacterial cultures were grown overnight in LB at 37°C, diluted 1:100 into 1.5 L LB, and grown (37°C, shaking) to OD_600_ 0.9-1.1. Cells were pelleted (10,000xg for 15 min 4°C), and supernatants were collected and concentrated to 100 mL using tangential flow with a 10 kDa MWCO filter (Pall). The retentate was then filtered (0.22 µm, Pall) to generate cell-free supernatant (CFS). To concentrate soluble proteins and OMVs from the CFS, ammonium sulfate was added to a final concentration of 90%, dithiothreitol added to 0.1 M, and the samples incubated overnight at 4°C with stirring. The precipitate was pelleted (15,000xg, 15 min, 4°C) and resuspended in 10 mL 20 mM HEPES, pH 8.0 (HEPES). The samples were then dialyzed in a ThermoFisher G2 cassette (10,000 MWCO) against HEPES, overnight at 4°C. The samples were further concentrated using Millipore 10,000 MWCO centrifugal filters, and the soluble proteins were separated from the OMVs using an iodixanol (Optiprep, Sigma) density gradient. Concentrated proteins and OMVs were adjusted to 50% Optiprep/HEPES in 2 mL, and applied to the bottom of the gradient tube, and subsequently layered with 2 mL of 40%, 2 mL of 35%, 4 mL of 30%, and 2 mL of 25% Optiprep/HEPES. The gradient was centrifuged overnight at 40,000xg, 4°C in an ultracentrifuge (SW 40 Ti swing bucket rotor, Beckman Coulter). Gradient fractions were removed from the top in 1 mL aliquots, and the lipid and protein content for each fraction were identified using FM4-64 and Bradford assays as described previously^29^. The OMV-containing fractions (typically 1-5) and soluble protein fractions (typically 7-12) were pooled and diluted with HEPES, then centrifuged at 40,000xg for 2 h to pellet OMVs and proteins and to remove Optiprep. Protein and OMV pellets were resuspended in HEPES, and sterile-filtered (0.4 µm PVDF, VWR).

### rPaAP purification

rPaAP expression, purification, and refolding were carried out as described previously^15^, and the refolded protein was concentrated using 10,000 MWCO Amicon filter units (Millipore-Sigma). The concentrated material was then dialyzed using the same method described above, applied to a S200 Sephadex ion exchange column, eluted with a gradient of 0-2 M NaCl, pH 7.6, at a flow rate of 0.02 mL/min, and 18- 2 mL fractions were collected. Fractions were assayed for leucine aminopeptidase activity as described^30^, and analyzed by SDS-PAGE and Ruby staining as described above. Fractions containing pure rPaAP were pooled and concentrated using Amicon 10,000 MWCO filter units to ∼0.5 µg/mL.

### Complementation and detachment experiments

For biofilm complementation assays, CFS, OMV fractions, soluble fractions, and rPaAP samples were prepared as described above, total protein was measured by Bradford assay, and aminopeptidase activity was measured as described above. For standard complementation experiments, samples were added to cocultures with new media at 1 hpi. 50 µg total protein was used for CFS samples, and 25 µg total protein was used for OMVs, soluble fractions, and rPaAP samples. For co-addition experiments, 25 µg of ΔPaAP OMVs were incubated with 25 µg of sPaAP or rPaAP for 10 min at room temperature prior to coculture treatment. In detachment experiments, 25 µg of OMV samples were added to cocultures at 4.5 hpi. For OMV pretreatment of A549 cell layers, 25 µg of OMV samples were added to A549 cells for 15 min prior to inoculation. For experiments matched by PaAP activity, samples were standardized to the aminopeptidase activity found in 25 µg (by total protein) of PaAP^+^ OMVs.

### Protease activity and inhibition

Protease activity was determined using the Pierce Protease Activity Kit (ThermoFisher Scientific). 10 µg of each OMV sample was used, and the rPaAP control was matched to the aminopeptidase activity found in this concentration of OMVs. Protease activity was inhibited by incubating the samples with 2X cOmplete Protease Inhibitor Cocktail (Millipore Sigma) and 10 mM phosphoramidon (Sigma) in 20 mM HEPES for 30 min at 37°C.

### Statistics

For all experiments, statistics were completed using GraphPad Prism t-tests unless otherwise indicated and the average +/- SEM is shown. For microscopy images and imaging quantifications, representative results are shown. Figures show a single biological replicate calculated from at least 5 technical replicate images. Images chosen for publication were the closest to the mean value for each experiment. For all other experiments, results are calculated from 3 biological replicates and the average +/- SEM is shown.

## Results

### PaAP expression limits *P. aeruginosa* early biofilm cellular mass and organization on host epithelial cells

Previous reports have described high expression of the *P. aeruginosa* aminopeptidase PaAP in both late-log phase and biofilm cultures, and PaAP expression was found to increase along with known virulence determinants when the bacteria are grown on lung epithelial cells^22^. To examine the hypothesis that PaAP impacts bacterial microcolony formation, we adapted a previously published biofilm coculture assay for use with A549 lung epithelial cells^26^. When grown over the course of several days, these cells form fully confluent cell layers that partially model the polarized epithelial monolayers that the bacteria colonize during infection. Once grown to confluency they can be inoculated with fluorescent bacteria and microcolonies can be assessed by CLSM.

To first establish the coculture conditions for our biofilm assays in which we would use a clinical *P. aeruginosa* isolate (S470 WT), the confluent A549 cells were inoculated with fluorescently labeled S470 and incubated for up to 6 h, and the A549 cells were assessed for damage and breaks in the cell layer. We established that this biotic substrate could be used to investigate clinical *P. aeruginosa* strain biofilm development through the early stages (up to 5 h post infection, hpi) of microcolony formation when LDH activity remained low (**Fig S1**) as substantial breakdown of these cellular layers occurred after 6h. We noted that once these cytotoxic bacteria infiltrated the host cell layer, the microcolony structure also broke down, as the bacteria were able to colonize the underlying tissue culture dish and migrate to the air-liquid interface. These results were consistent with the previously described toxicity of *P. aeruginosa* clinical strains to host cells^31^ and time courses for static coculture biofilm models^26^. In addition, we confirmed that the bacterial colonies exhibited biofilm properties, as they displayed gene expression patterns similar to those documented for biofilm-associated cells (**Fig S2**) as well as increased antibiotic resistance when compared to planktonically grown bacteria (**Fig S3**).

To examine the effect of PaAP on *P. aeruginosa* biofilm microcolony growth, we quantitatively compared cocultures of fluorescent S470 WT to S470 ΔPaAP, a previously described isogenic knockout strain (S470 ΔPaAP) that contains a disruption of PA2939, the gene encoding PaAP^24^. We observed significant differences between the two biotic biofilms at 5 hpi. Interestingly, the PaAP mutant strain formed biofilms with substantially greater cellular biomass compared to the WT strain (Fig 1A, B). We also examined S470 WT and ΔPaAP microcolony morphologies (Fig 1D, E). In addition to their greater quantitative biomass, S470 ΔPaAP colonies displayed greater cellular organization. Even at this early point in biofilm development, characteristic mushroom-shaped colony structures were observed for the PaAP mutant. The S470 WT microcolonies, by comparison, showed very little cellular organization and were packed less densely. Based on these data, we can conclude that the lack of PaAP not only increases cellular biomass, but also cellular organization at the microcolony level during early coculture biofilm development.

**Figure 1:**
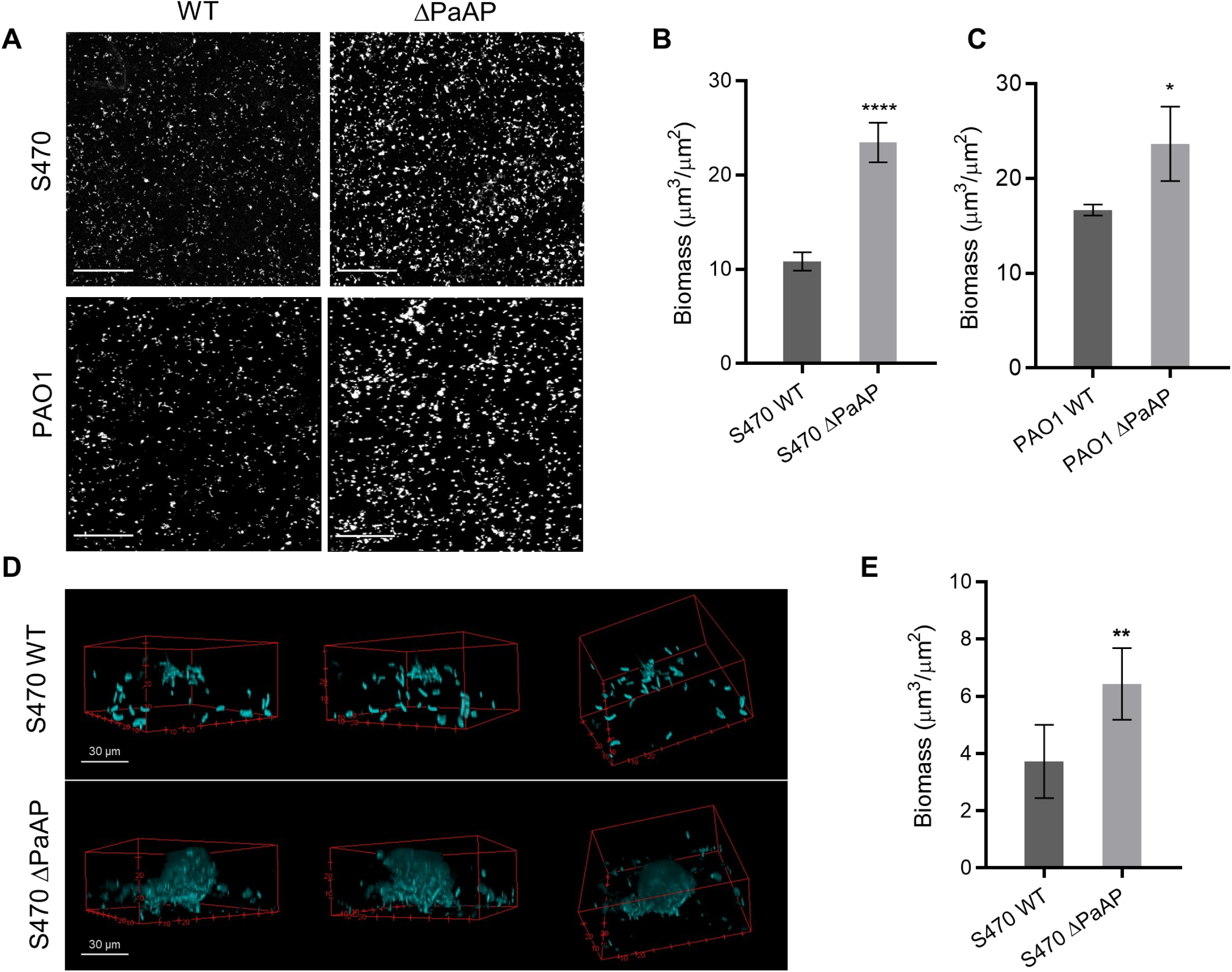
Deletion of the PaAP aminopeptidase increased the density, biomass, and organization of bacterial biofilms on host cells. **(A)** S470 and PAO1 WT and ΔPaAP strains were inoculated onto confluent A549 cell layers, and images were taken at 10X magnification, 5 hpi and **(B, C)** the images were quantified. **(D)** For cocultures as described for **A**, images of microcolonies were taken using 100X magnification at 5 hpi and **(E)** quantified. For all experiments, representative results are shown. Quantified bacterial biomass in each set of cocultures was compared; *p<0.05, **p<0.01, ***p<0.001, ****p<.0001. Scale bars: 200 µm, unless otherwise specified.

To confirm this PaAP-dependent phenotype in a different strain background, we compared the biofilm characteristics of PAO1, a laboratory *P. aeruginosa* strain commonly studied *in vitro,* and the isogenic PAO1 ΔPaAP strain that harbors a transposon insertion in PA2939. We note that our lab previously showed that OMVs isolated from PAO1 cultures show very low aminopeptidase levels compared to S470 and other clinical *P. aeruginosa* isolates^13^. Nevertheless, as with the clinical strain phenotype, PAO1 ΔPaAP formed significantly more biomass as compared to the WT counterpart (Fig 1A, C). Our observations indicate that PaAP is important to biofilm development, even in strains with low endogenous PaAP expression, and that its role may be relevant to a broad range of *P. aeruginosa* strains.

### Increased biofilm cellular density induced by PaAP deletion coincides with PaAP expression and increased EPS production

To detail the time course of the PaAP-dependent phenotype, we imaged the cocultures at hourly intervals post infection. Directly following the wash step at 1 hpi and up to 3 hpi, we observed a similar level of biomass from the S470 WT and ΔPaAP strains. A significant phenotype did not develop until 4 hpi (Fig 2A), suggesting that the aminopeptidase is critical in limiting microcolony growth and development rather than during the initial steps of colonization and attachment. Based on this timeline, it is expected that the observed phenotype would develop in parallel with the level of aminopeptidase expression. Indeed, previous reports found PaAP to be regulated by the *las* quorum sensing system^14, 32^, and thus PaAP expression would be expected to increase as bacterial density increases in our experimental system. Using qRT-PCR, we examined the expression of PaAP in S470 WT biofilms over time under the coculture assay conditions. Consistent with quorum sensing density-dependent expression, our results show expression of PaAP mRNA at 1 hpi, increasing steadily over the next several hours (**Fig S4**) and PaAP protein in cocultures at the 5 hpi time point (Fig 2B). Based on these data, it appears that the biofilm limiting phenotype may depend on either a threshold level of aminopeptidase or a specific microcolony developmental phase.

**Figure 2:**
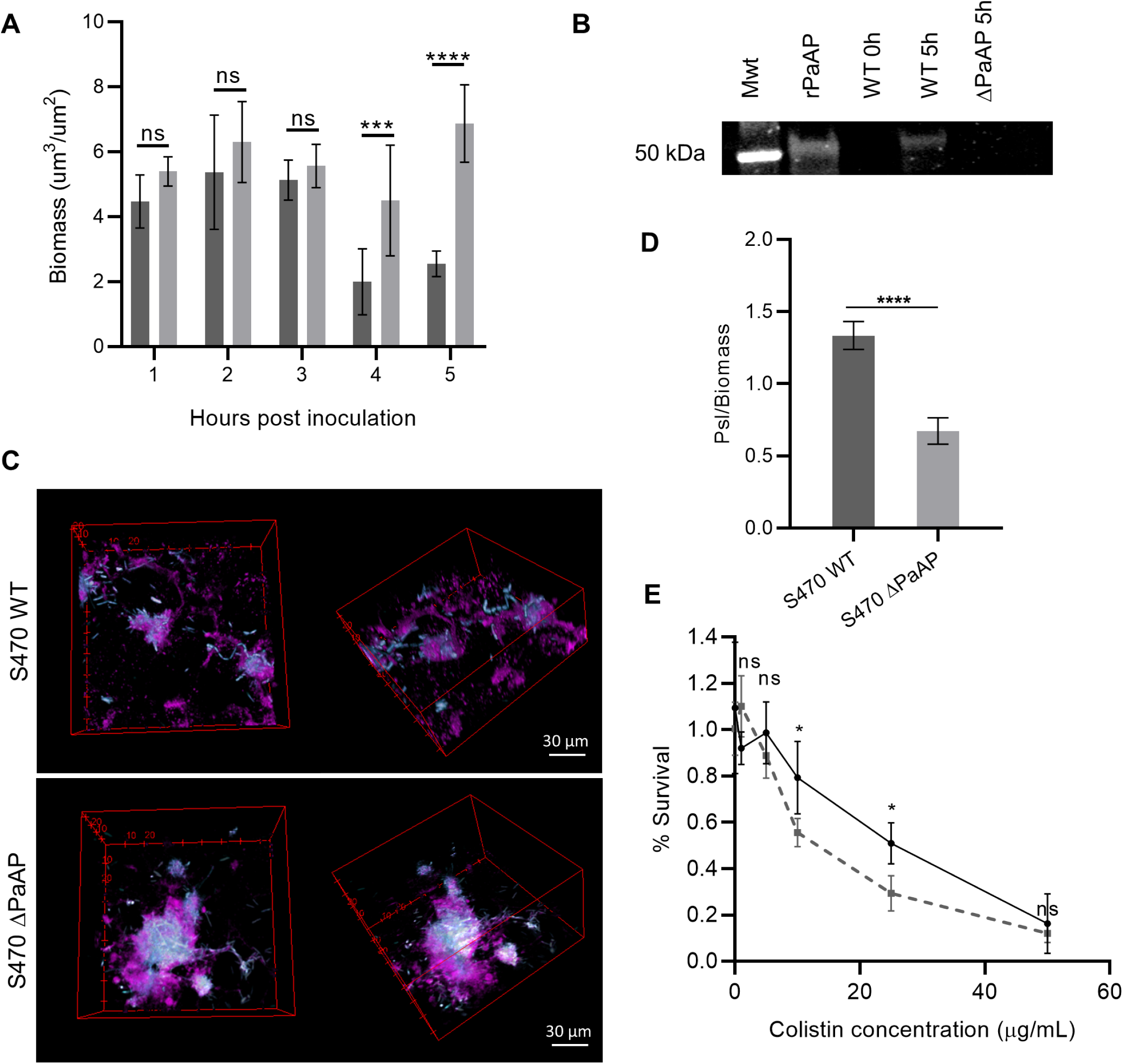
PaAP expression promotes Psl production in coculture biofilms. **(A)** Bacterial biofilms cocultured on A549 cells were imaged by confocal microscopy at indicated time points. (S470 WT: dark gray, S470 ΔPaAP: light gray). **(B)** Total protein was extracted from S470 WT and ΔPaAP cocultures at indicated hpi and precipitated, and PaAP was detected in the samples using Western blot. Mwt, 50 kDa standard; PaAP, purified PaAP control. **(C)** Bacterial biofilms (cyan) were cocultured on A549 cells for 5 h, stained with HHA-FITC lectin (magenta) and imaged by confocal microscopy and **(D)** the ratio of Psl/cellular biomass was calculated. **(E)** At 3 hpi, S470 WT (solid line) and ΔPaAP (dashed line) cocultures were treated with the indicated concentrations of colistin. Survival indicates the percent of biomass remaining attached to the host cell substrate. For all experiments, representative results are shown. *p<0.05, **p<0.01, ***p<0.001, ****p<.0001, ns, not significant.

Bacterial fluorescence images from the coculture assay in Fig 1 revealed significant density and structural differences between the biofilms formed by WT and ΔPaAP *P. aeruginosa* strains, but we were also interested in whether PaAP affected the composition of the biofilm matrix. The *P. aeruginosa* matrix is composed of polysaccharides, proteins, DNA, and vesicles secreted by cells in the microcolony. We imaged and quantitated the polysaccharide content of the matrix using a lectin specific for Psl, a major component of non-mucoid matrices. As seen in Fig 2C and 2D, the S470 microcolonies contained significantly more Psl per cellular biomass. This also correlated with a modest, but significant increase in colistin resistance (Fig 2E), suggesting that the aminopeptidase-dependent modulation of matrix can result in an overall more robust biofilm structure.

### Leucine aminopeptidase expression mediates pellicle growth under specific nutrient conditions

While our coculture model has provided strong evidence that PaAP affects the growth and matrix development of biofilm microcolonies grown on a host epithelial surface, *P. aeruginosa* is also known to form other biofilm structures. When grown in liquid media, *P. aeruginosa* forms biofilm communities called pellicles at the air-liquid interface. During pellicle formation, the bacteria attach to the culture container and biofilm development can be quantitatively assessed using a Crystal Violet stain^28^ or live-dead staining of the pellicle bacteria^33^.

Recently, Zhao et al showed that PaAP can affect PAO1 *P. aeruginosa* biofilm pellicles formed in cultures using Jensen’s media at 30°C^33^. While the body of this work mainly focused on a late-stage phenotype, they also observed that PaAP deletion led to increased biomass in early biofilms. By growing the bacteria planktonically in Jensen’s media to mid-log phase and diluting and inoculating the culture containers, we were able to replicate this phenotype with the S470 clinical strain and corresponding PaAP deletion mutant (Fig 3A, B, C**, Untreated**). In Jensen’s media, the ΔPaAP strain formed pellicles with significantly greater cellular biomass. Interestingly, this phenotype was not found with cultures grown in other media used frequently in biofilm studies (M63 minimal media and LB + glycerol) or the cell culture media used in the coculture experiments (both fresh and conditioned F-12K) (**Fig S5**). These data lead us to conclude that PaAP-dependent effects in early biofilms are growth-condition and nutrient specific.

**Figure 3:**
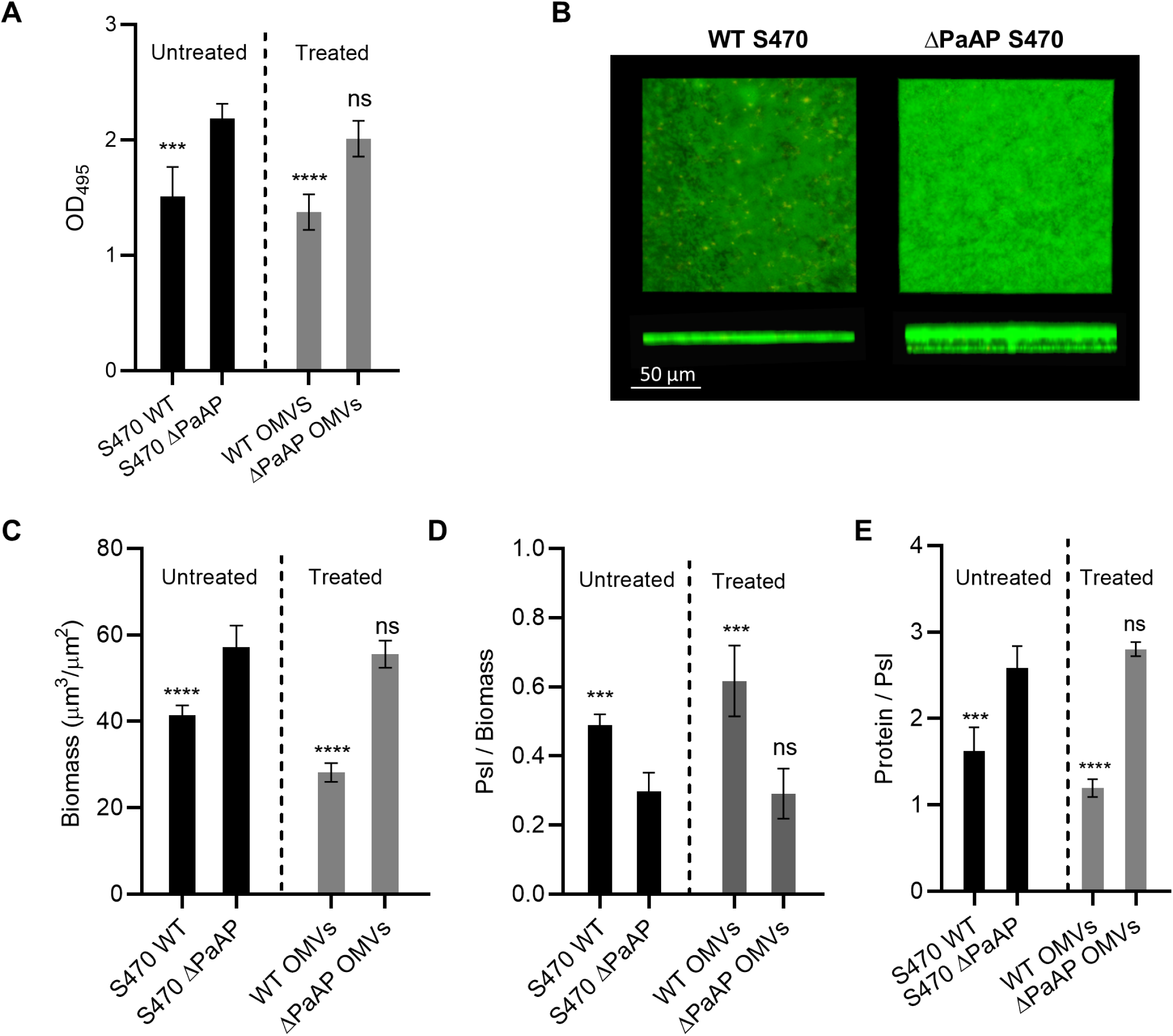
PaAP deletion modifies pellicle cellular biomass and matrix composition and WT OMV treatment complements these phenotypes. **Untreated: (A)** S470 WT or ΔPaAP were grown in 35mm dishes for 6 h and pellicle biofilms were quantified by Crystal Violet staining (OD_495_). **(B)** Pellicles formed by S470 WT or ΔPaAP after 6 h of growth were live/dead stained and imaged by confocal microscopy **(C)** and their biomass calculated. Pellicles were stained for **(D)** matrix Psl and **(E)** total protein, and the results quantified and normalized to cellular volume and Psl volume, respectively. **Treated: (A, C-E)** ΔPaAP pellicles were treated at 1 hpi with S470 WT or ΔPaAP OMVs and imaged at 5 hpi. Untreated WT and treated ΔPaAP pellicles were stained as above, quantified, and values compared to untreated ΔPaAP pellicles; *p<0.05, **p<0.01, ***p<0.001, ****p<.0001, ns, not significant. For microscopy experiments, representative results are shown.

To examine whether PaAP affected the matrix composition of abiotic biofilms, pellicles grown in Jensen’s media were stained for Psl. As with the coculture biofilms, the S470 WT pellicles consisted of significantly greater exopolysaccharide per cellular biomass as compared to ΔPaAP pellicles (Fig 3D, Untreated). We were also able to stain for matrix protein content using this model since the background protein levels were much lower as compared to the coculture biofilms. Compared to the ΔPaAP strain, the WT pellicles exhibited a significantly lower protein to Psl ratio (Fig 3E, Untreated), strengthening our conclusion that PaAP expression can cause substantial changes in matrix composition of early biofilms under the stated growth conditions.

### PaAP^+^ OMVs, but not soluble aminopeptidase, exhibit anti-biofilm activity

Complementation of the PaAP deletion phenotype was pursued next. Based on its tightly-controlled, quorum sensing-regulated expression and secretion pathways, PA2939 complementation using plasmid-based expression has proven to be unfeasible. We therefore performed biochemical complementation experiments to test the ability of secreted PaAP to reconstitute the original biofilm composition in ΔPaAP biofilms. Complementation was tested using cell-free supernatants (CFS) from mid-log phase cultures of S470 WT and S470 ΔPaAP after confirming the respective presence and absence of PaAP activity in the S470 WT and S470 ΔPaAP CFS by Western blot (Fig 4A) and aminopeptidase activity assays (**Fig S6**). Equivalent amounts (by total protein) of CFS were added to washed S470 ΔPaAP coculture biofilms 1 hpi, and the cocultures imaged 5 hpi. Addition of the PaAP-containing S470 WT CFS complemented the phenotype of ΔPaAP biofilms to WT biomass levels, while the S470 ΔPaAP CFS resulted in an insignificant change (Fig 4B). We concluded that *P. aeruginosa* CFS harbors a PaAP-dependent biofilm modulating activity.

**Figure 4:**
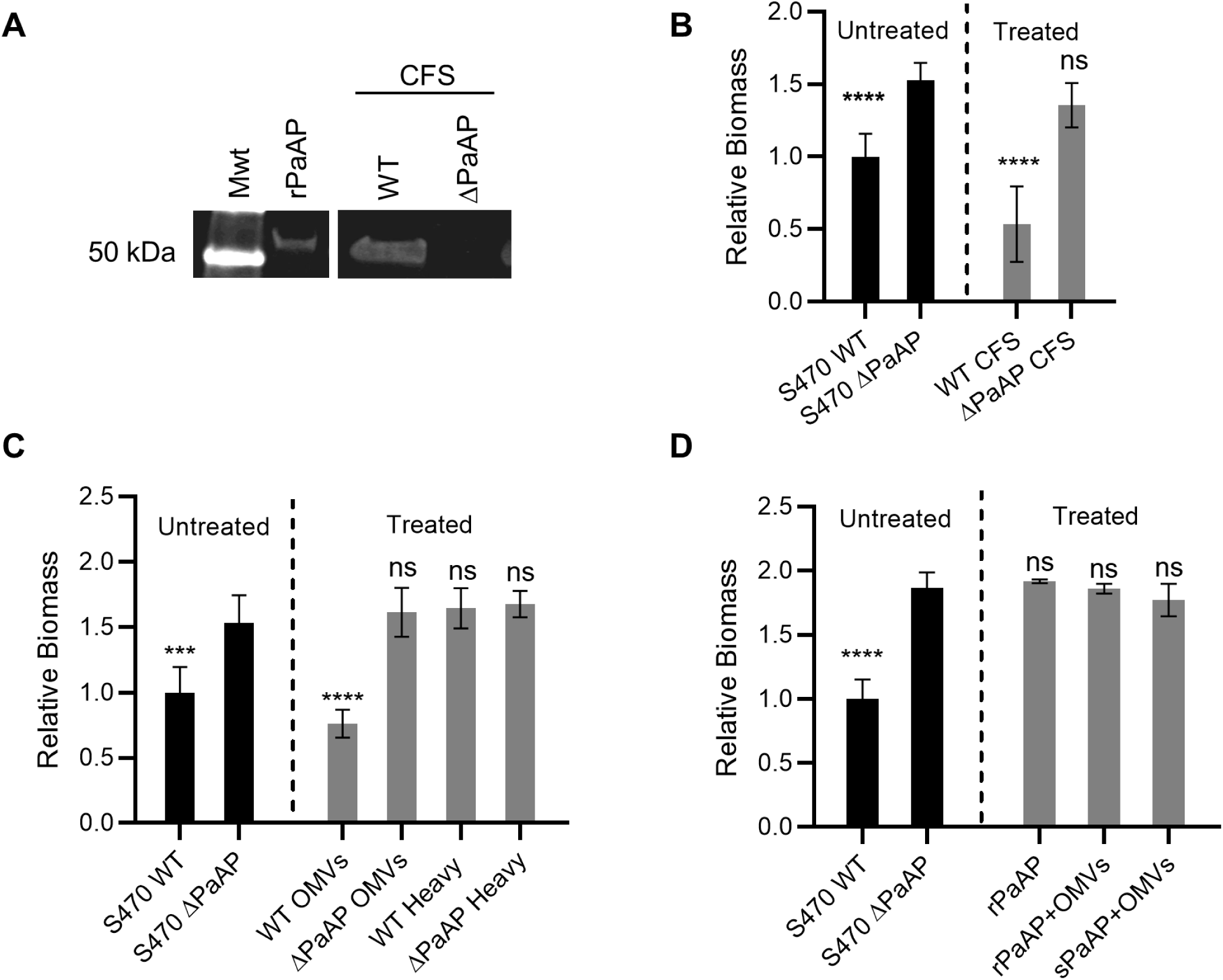
Addition of PaAP^+^ OMVs inhibits formation of *P. aeruginosa* coculture biofilms. **(A)** Cell free supernatants (CFS) from S470 WT and S470 ΔPaAP cultures were TCA-precipitated, and PaAP was detected by immunoblotting after separation of samples by SDS-PAGE. The relative migration of rPaAP and the 50 kDa molecular weight standard (Mwt) are shown. Samples were run on the same gel, and an intervening lane was excised in this figure. S470 ΔPaAP biofilms cocultured with A549 cells were treated with CFS S470 WT or ΔPaAP at 1 hpi **(B)**, with fractions containing either light (OMVs) or dense (heavy) from S470 WT or S470 ΔPaAP CFS **(C)**, or with rPaAP, rPaAP with ΔPaAP OMVs, or sPaAP with ΔPaAP OMVs **(D)**. The biomass of these treated biofilms and the S470 WT untreated control were quantified at 5 hpi and were compared with untreated S470 ΔPaAP biofilms; *p<0.05, **p<0.01, ***p<0.001, ****p<.0001, ns, not significant. For all experiments, representative results are shown.

We wondered if secreted PaAP itself was responsible for the activity and recalled that after secretion by the type II secretory system, PaAP is found both associated with OMVs and as a soluble protein^24^. Therefore, to locate the source of the activity in the coculture supernatants, soluble proteins and OMV-bound factors were separated using iodixanol density gradients, and consistent with our previous reports^13^, aminopeptidase activity was detected in both the light- and heavy-density fractions, representing the OMV-bound and soluble forms of PaAP (sPaAP), respectively. Notably, whereas treatment with the OMV-containing S470 WT fractions was able to significantly inhibit cellular biomass development by S470 ΔPaAP and partially restore relative polysaccharide levels in these biofilms, the heavier, sPaAP-containing fractions had no noticeable effect on biomass (Fig 4C, **Fig S7**). As expected, neither treatment at 1 hpi of OMV-containing or heavy fractions of the CFS from S470 ΔPaAP cultures altered the ΔPaAP coculture biomass at 5 hpi (Fig 4C). PaAP^+^ OMVs, but not ΔPaAP OMVs, were also able to complement the biofilm phenotype observed in Jensen’s media pellicles and could restore matrix composition in these samples (Fig 3A, C-E, Treated). These data suggested OMVs were responsible for the biofilm modulating activity in a PaAP-dependent manner, and that solubly secreted PaAP was not.

To confirm that aminopeptidase alone was not capable of restoring these phenotypes, we purified and refolded recombinant PaAP (rPaAP) expressed in *E. coli*^15^. Consistent with the results using sPaAP, purified rPaAP was also incapable of disrupting the formation of coculture biofilms by the ΔPaAP strain (Fig 4D). Finally, we tested whether addition of rPaAP or sPaAP could reconstitute the biofilm inhibitory activity of the OMVs isolated from ΔPaAP cultures (ΔPaAP OMVs). These rPaAP/ΔPaAP OMV and sPaAP/ΔPaAP OMV preparations also failed to inhibit biotic biofilm development by the S470 ΔPaAP strain (Fig 4D). These data lead us to conclude that OMVs derived from a PaAP^+^ strain, mediate cell aggregation and matrix composition at early stages of biofilm microcolony development. The fact that ΔPaAP OMVs and PaAP alone or in combination with ΔPaAP OMVs were unable to limit biofilm development additionally suggests that the aminopeptidase is not directly causing changes in biofilm architecture, but rather modifying OMV activity and thereby indirectly changing the biofilm microcolonies. ^24^

### PaAP^+^ OMVs exhibit biofilm detachment activity against non-self *P.* aeruginosa and *K. pneumoniae*

In order to characterize the biofilm-remodeling mechanism of the OMVs, we studied the kinetics of the vesicle-mediated anti-biofilm activity. OMVs were added to cocultures either before inoculation (pre-treatment), at 1 hpi, or at 4.5 hpi, (0.5 h before the final imaging time point). Table 1 summarizes the findings of these experiments. Differences between the WT and ΔPaAP OMV pre-treated cocultures did not develop until the late time point, providing evidence that the WT OMVs did not influence the ability of bacteria to attach to the host cell surface. Furthermore, WT OMV treatments 1 hpi did not substantially reduce microcolony structures until 3 h and 5 h post-infection, consistent with an effect by OMVs on microcolony development rather than attachment.

**Table 1:** PaAP^+^ OMVs inhibit microcolony development and disrupt pre-formed microcolonies

| OMV Addition<sup>1</sup> | Imaging<sup>2</sup> | OMVs | Biomass<sup>3</sup> | Interpretation |
| --- | --- | --- | --- | --- |
| Bacterial Attachment |  |  |  |  |
| pre-treatment | 1 hpi | None (control) | = | OMV pretreatment of the A549 cells does not affect initial bacterial cell attachment to the host surface.<br>This suggests the mechanism behind PaAP<sup>+</sup> OMV biofilm remodeling is not related to attachment |
|  |  | S470 WT | = |  |
|  |  | S470 ΔPaAP | = |  |
| Microcolony Inhibition |  |  |  |  |
| pre-treatment | 5 hpi | None (control) | = | Addition of WT OMVs but not ΔPaAP OMVs at any point early in the coculture experiment results in lower bacterial biomass.<br>This effect is more pronounced at later time points, suggesting OMV-mediated biofilm inhibition acts on a bacterial pathway important to microcolony growth and formation, rather than initial attachment. |
|  |  | S470 WT | - - |  |
|  |  | S470 ΔPaAP | = |  |
| 1 hpi | 3 hpi | None (control) | = |  |
|  |  | S470 WT | - |  |
|  |  | S470 ΔPaAP | = |  |
| 1 hpi | 5 hpi | None (control) | = |  |
|  |  | S470 WT | - - |  |
|  |  | S470 ΔPaAP | = |  |
| Microcolony Detachment |  |  |  |  |
| 4.5 hpi | 5 hpi | None (control) | = | WT OMVs, but not ΔPaAP OMVs, can disrupt established microcolonies, further suggesting that the OMVs affect microcolony growth and cell release mechanisms. |
|  |  | S470 WT | - - - |  |
|  |  | S470 ΔPaAP | = |  |

<sup>1</sup>S470 ΔPaAP cocultures were treated with the indicated OMVs at the indicated times. For pre-treatment, A549 cells were incubated with vesicles for 0.5 h before bacterial inoculation.
<sup>2</sup>Samples were imaged at the indicated times and assessed for biomass.
<sup>3</sup>Biomass is represented relative to the untreated S470 ΔPaAP controls for each experiment (=: no significant difference; +: significant increase; -: significant decrease). The number of +/- quantifies the relative biomass differences (+/-: 5-25%; +/- -: 25-50%; +++/- -: >50%).

To test whether PaAP^+^ vesicles could disrupt preformed microcolony structures, OMVs were not added to the A549/S470 ΔPaAP cocultures until 4.5 hpi, and the biofilms imaged 30 minutes later. As seen in Fig 5A and B, this brief PaAP^+^ OMV treatment resulted in a significant and very substantial decrease in biomass of a preformed biofilm. To examine whether this biofilm reducing activity was specific to S470 strain-derived vesicles, we purified and tested OMVs derived from the PAO1 and PA14 *P. aeruginosa* WT strains and their corresponding ΔPaAP mutants. WT OMVs from both strains disrupted the preformed S470 ΔPaAP coculture biofilms whereas OMVs from the isogenic mutants did not (Fig 5A, B). OMVs also mediated biofilm reduction for S470 pellicles grown in Jensen’s media but not for pellicles that did not show the original phenotype (e.g. those grown in M63 minimal media or tissue culture cell media) (Fig S8).

**Figure 5:**
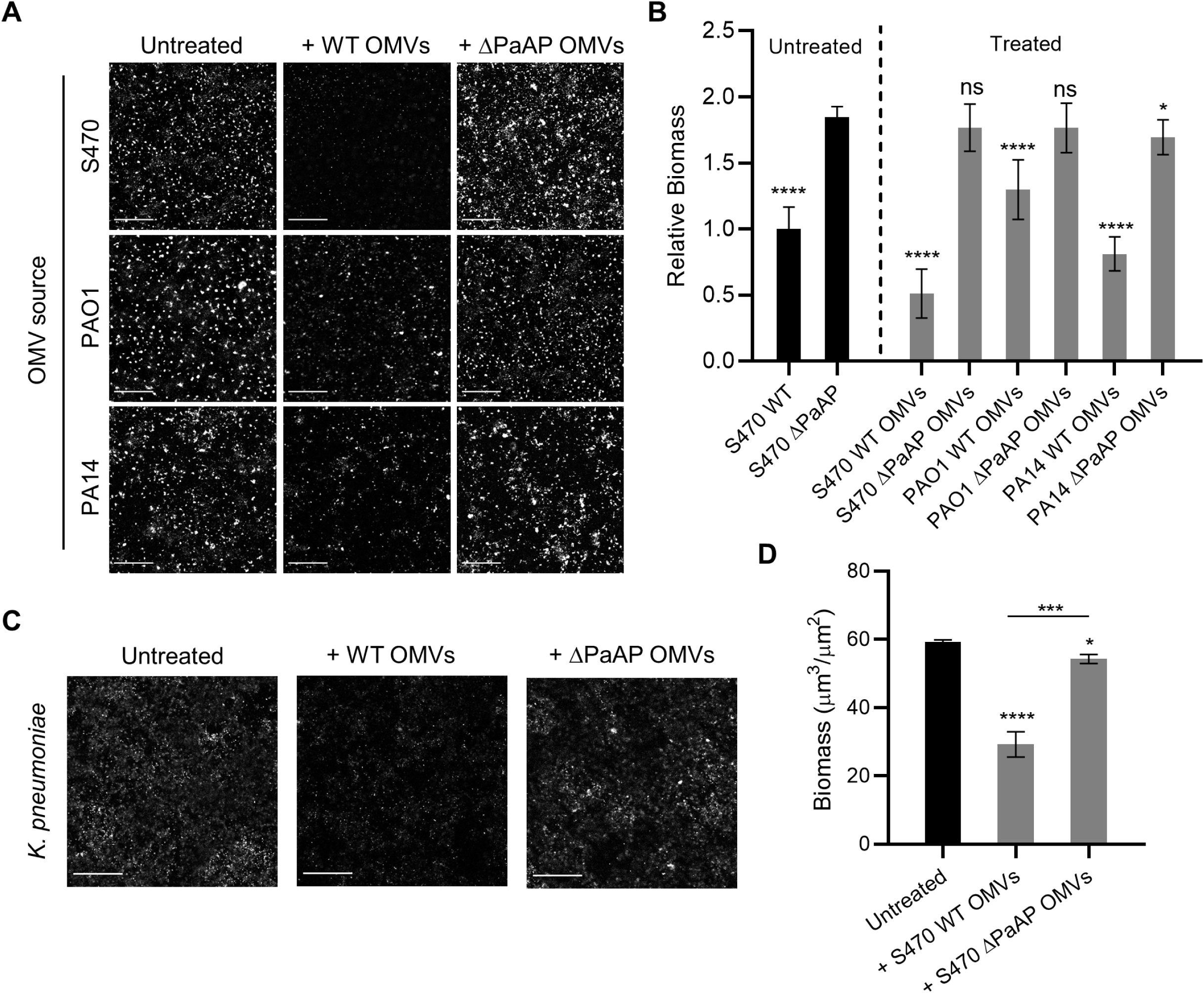
PaAP^+^ OMVs cause biofilm detachment in both pseudomonads and non-pseudomonads. **(A)** Top row: OMVs from WT or ΔPaAP cultures of the indicated *P. aeruginosa* strains were added to S470 ΔPaAP biofilm cocultures at 4.5 hpi, 30 minutes prior to imaging and **(B)** these results were quantified. The biomass of treated biofilms and the S470 WT untreated controls were quantified and were compared with untreated S470 ΔPaAP biofilms**. (C)** *K. pneumoniae* cocultures were treated with S470 WT or ΔPaAP vesicles at 4.5 hpi, biofilm formation was assessed by Congo Red staining at 5 hpi, and **(D)** quantified. The biomass of treated *K. pneumoniae* biofilms were quantified and were compared with untreated biofilms. For all experiments, representative results are shown. *p<0.05, **p<0.01, ***p<0.001, ****p<.0001, ns, not significant. Scale bars: 200 µm.

During respiratory infections biofilms are often found as polymicrobial communities, and components secreted by one species can impact the microcolonies of other strains or species in nearby environments. To determine whether the anti-biofilm activity of the OMVs is species-specific, A549 cell layers were inoculated with *K. pneumoniae*, a bacterium which is often found with *P. aeruginosa* in cases of ventilator-associated pneumonia^34^. S470 WT or ΔPaAP OMVs were added at 4.5 hpi, and the cocultures were stained at 5 h with Congo Red to examine both cellular and matrix biomass. As seen in (Fig 5C), S470 WT vesicles significantly reduced *K. pneumoniae* total biomass, while ΔPaAP OMVs had very little effect. These data suggest that the OMVs target a conserved feature of biofilm architecture, allowing them to impact the colonization ability of multiple clinically relevant pathogens.

### Protease inhibitors decrease biofilm detachment activity of *P. aeruginosa* vesicles

Antimicrobial resistance is often used to inform whether cells released from biofilms retain characteristics of the parent biofilm or have transitioned to a planktonic state. Previous studies have shown that mechanical detachment of biofilms does not restore antimicrobial susceptibility; however, cells undergoing a signaled dispersal response will lose this resistance as they gain motility and release from the biofilm colonies. To characterize which of these occurs during vesicle-mediated cellular release from coculture biofilms, the released cells were collected and assayed for colistin resistance (Fig 6A). These cells displayed significantly higher antibiotic resistance compared to planktonically grown cultures, suggesting that the mechanism behind OMV-mediated biofilm disruption and remodeling likely involves the detachment of cells or cellular aggregates, rather than a programmed dispersal response.

**Figure 6:**
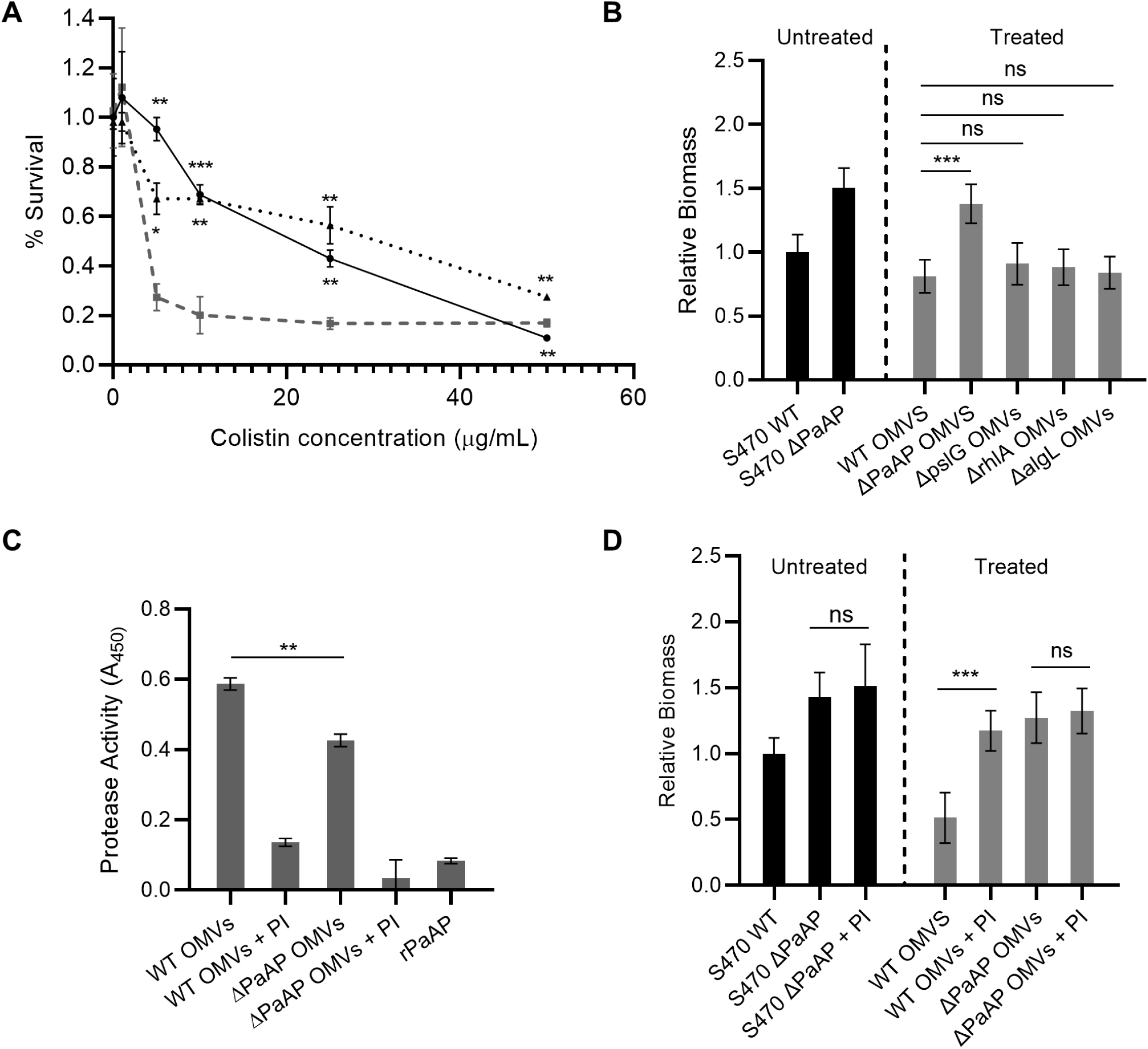
PaAP increases protease activity in OMVs, leading to cell detachment from biofilm microcolonies. **(A)** S470 ΔPaAP cocultures (solid line) were challenged with colistin at 3 hpi and examined for remaining biomass at 5 hpi. Additional S470 ΔPaAP cocultures were treated with S470 WT OMVs at 4.5 hpi, the detached cells (dotted line) were collected at 5 hpi and challenged with colistin for 2 h. These samples were compared to planktonically grown S470 ΔPaAP cells (dashed line) challenged with colistin at 3 hpi with live/dead assessment at 5 hpi. **(B)** S470 ΔPaAP biofilm cocultures were treated with OMVs isolated from PA14 WT and the indicated PA14 transposon insertion mutants at 4.5 hpi, and the biomass was quantified at 5 hpi. The biomass of treated biofilms and the S470 WT untreated control were quantified, and treated samples were compared to biofilms treated with S470 WT OMVs**. (C)** Protease activity was measured in the indicated S470 WT and ΔPaAP OMV preparations with and without protease inhibitor cocktail (PI) and for rPaAP. **(D)** S470 ΔPaAP biofilm cocultures were treated with the indicated S470 WT and ΔPaAP OMVs with and without PI at 4.5 hpi and the resulting biomass was quantified at 5 hpi. The biomass of treated biofilms, the S470 WT untreated control, and the S470 ΔPaAP biofilm incubated with PI were quantified and samples were compared to the corresponding samples without PI. For microscopy-based experiments, representative results are shown. For all other experiments, n=3. *p<0.05, **p<0.01, ***p<0.001, ****p<.0001, ns, not significant.

Based on the evidence provided in Fig 4, this cellular detachment is likely not caused directly by the aminopeptidase, but instead relies on an indirect effect of PaAP on OMV cargo. Several factors from *P. aeruginosa* are known to facilitate biofilm detachment directly, including those that can break apart the biofilm structure by enzymatic or biochemical means. We studied the impact of three known disruption factors on this phenotype: rhamnolipids^35, 36^, and the exopolysaccharide hydrolases PslG^33, 36^ and AlgL^36^. Vesicles were isolated from cultures of PA14 transposon mutants with a deletion in each potential target and evaluated for their ability to reduce the biomass of 4.5 hpi S470ΔPaAP coculture biofilms. As shown in Fig 6B, none of these mutations significantly changed the detachment activity observed for PA14 WT vesicles.

Known biofilm-detachment factors include not only polysaccharide hydrolases, but also DNase, Proteinase K and various endogenous *P. aeruginosa* proteases^36^. Based on the importance of endogenous proteases to biofilm development, we tested PaAP^+^ and ΔPaAP OMVs for overall protease activity. Interestingly, we found that PaAP^+^ OMVs display higher protease activity, which cannot be directly accounted for by the presence of aminopeptidase in these samples, as demonstrated by the inclusion of an rPaAP control matched to aminopeptidase activity levels in the PaAP^+^ vesicles (Fig 6C). By pre-incubating the vesicle samples with a protease inhibitor (PI) cocktail, we were able to eliminate both protease and anti-biofilm activity in the PaAP^+^ OMVs (Fig 6C, D) without impacting aminopeptidase activity in the samples (**Fig S9**). While future work will be required to fully elucidate the proteases involved in this phenotype, these results suggest that PaAP expression in *P. aeruginosa* increases endogenous protease activity that becomes localized to secreted OMVs, which can in turn lead to OMV-induced cell detachment and contribute to remodeling of overall biofilm architecture.

## Discussion

We set out to investigate the role of the PaAP leucine aminopeptidase in biofilm development, as PaAP is one of the most abundant proteins in the *P. aeruginosa* biofilm extracellular matrix and one of the major protein components of OMVs from clinical *P. aeruginosa* isolates^13, 20^. When we compared coculture biofilms of wild-type *P. aeruginosa* and a PaAP deletion mutant, we found the mutant developed early biofilms with a significantly higher cellular biomass and a lower concentration of the matrix exopolysaccharide Psl, a phenotype confirmed in specific pellicle biofilm growth conditions. The mutant phenotype included a less robust biofilm architecture that rendered the cellular community more susceptible to the antibiotic colistin. We further found that biofilm detachment activity resided in OMVs in supernatant fractions from wildtype cultures, but not OMVs secreted by ΔPaAP cells. The effect of PaAP expression was indirect, as neither recombinant nor purified soluble aminopeptidase alone or in combination with ΔPaAP OMVs possessed detachment activity. Our data further lead us to conclude that PaAP influences changes in biofilm architecture through proteolytic activity, as WT OMVs exhibited higher proteolytic activity than ΔPaAP OMVs, and WT OMV-mediated detachment was sensitive to protease inhibition. OMVs have been implicated in the process of biofilm formation^7, 19^, but while their presence in these bacterial communities has been documented, active roles for these particles and the bacterial products they carry have not been extensively studied. Recently, Zarnowski et al have reported a role for vesicles in the biogenesis of antimicrobial-resistant *Candida albicans* biofilm matrices and have provided evidence that vesicle cargo proteins are “functional passengers” that can affect biofilm structure^37^. Similarly, our results suggest that vesicle-associated enzymes functionally impact *P. aeruginosa* biofilms.

Detachment of cells form the biofilm structure is a process previously shown to be able to be mediated by a variety of enzymes and biosurfactant molecules native to *P. aeruginosa*. Polysaccharide hydrolases, including both PslG and alginate lyase, have been described by multiple studies as mediating the release of cells from biofilm structures^36^. Zhao et al reported on the involvement of PslG in biofilm disruption, which was also, intriguingly, PaAP-dependent^33^. However, in this case, biofilm disruption was found to take place in late log-stage biofilms under nutrient-limiting conditions such that the lack of PaAP lead to cell lysis, releasing the periplasmic PslG enzyme. By contrast, the biofilm phenotype we observed not only does not depend on starvation, it is mediated by OMVs and is independent of PslG expression. Additionally, AlgL and rhamnolipids, which have biosurfactant properties that can cause the release of biofilm cells, were shown not to be involved in this effect. These results demonstrate a distinct role for both PaAP and OMVs in the modulation of biofilm communities.

Other described mechanisms to influence bacterial detachment involve protease activity, which is important in many aspects of *P. aeruginosa* biology^38^. Previous studies have implicated endogenous proteases in biofilm detachment and the dissolution of cellular aggregates^39^, specifically those with a protein rich matrix. Based on our data we can add new details to this model: endogenous PaAP expression helps regulate protease activity in secreted vesicles which, in turn, lead to the remodeling of biofilm architecture (Fig 7). The changes induced by OMVs ultimately increase Psl/biomass ratios in early biofilm matrices and help to protect the developing colony from disruption by antimicrobials. As for the specific protease(s) and substrate(s) involved in his mechanism, protease expression and activation during *P. aeruginosa* development is a highly complex, regulated process with many steps, including expression, secretion, and processing. Additionally, many of these proteases are involved in the activation and expression of others^15, 38, 40^. This complex system makes it challenging to specify which protease(s) are affected by PaAP expression, whether this is the effect of one or several endogenous proteases, and how PaAP increases vesicle-associated protease activity. The protein substrate(s) for the OMV-activity are also unknown and also require further study, however one interesting observation is that both early biofilms of *P. aeruginosa* and *K. pneumonia* contain a substrate for the activity.

**Figure 7:**
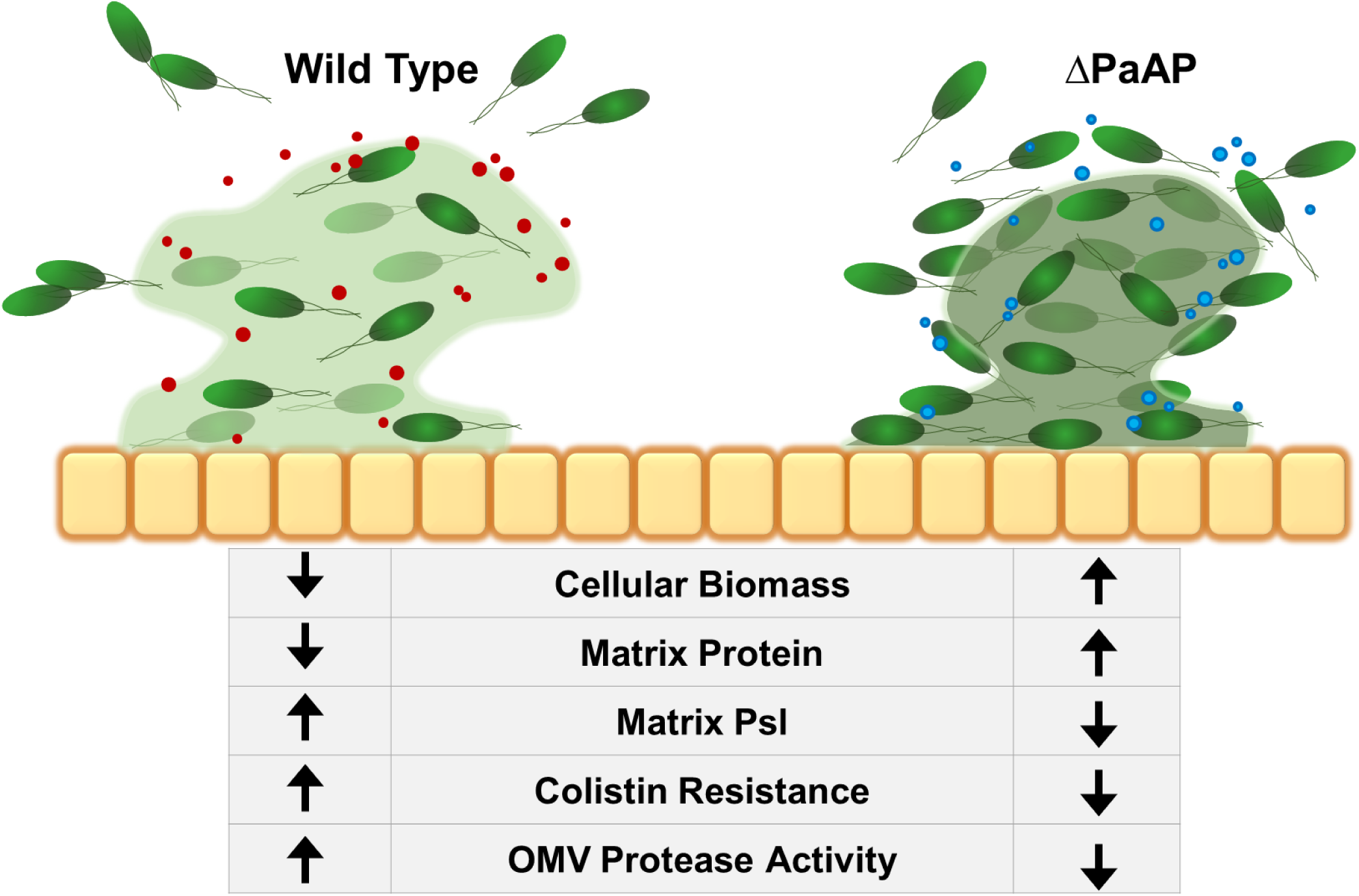
Summary of the effect of PaAP^+^ OMVs on early biofilm development. During WT growth, bacterial microcolonies secrete PaAP^+^ OMVs with increased endogenous protease activity. This leads to increased cell detachment from the colony structure and allows for increased matrix Psl polysaccharide production and protection for the cells against antibiotic treatment. Orange squares: A549 epithelial cells; Green cells: *P. aeruginosa* cells; Red circles: PaAP^+^ OMVs; Blue circles: ΔPaAP OMVs; Green haze: matrix Psl. Darker matrix color indicates a higher concentration of protein.

While some aminopeptidases have been implicated in pathogenesis phenotypes^41^, these enzymes are generally thought to act during starvation responses, in which they can provide necessary amino acids for cellular processes. This study, by contrast, presents a novel role for a bacterial aminopeptidase, as PaAP impacts early biofilm formation processes early in development and under nutrient-rich conditions. Additionally, because we have shown that this phenotype is not directly caused by the vesicle-associated aminopeptidase, but rather other vesicle-associated proteases, we have implicated PaAP in a more complex regulatory network that influences the activity of bacterial molecules and structures important to the formation of robust biofilm structures. Future work on this project will seek to elucidate the mechanism by which PaAP influences OMV-mediated biofilm maturation.

## Acknowledgements

We thank David FitzGerald (National Cancer Institute, Bethesda, MD) for anti-PaAP antibody and rPaAP constructs, and acknowledge the contribution of the dTomato plasmid by Carol Kim (University of Maine, Orno ME).

---

**Table S1: qRT-PCR primers used in this study**

**Figure S1: Biofilm coculture LDH release assay**

Supernatant samples were taken from biofilm cocultures at the indicated timepoints and tested for LDH release. S470 WT: dark gray; S470 ΔPaAP: light gray. Max indicates maximum LDH release for the samples.

**Figure S2: Coculture S470 biofilm gene expression reflects known biofilm patterns**

RNA was isolated from planktonic cells, pellicle biofilms, and coculture biofilms, and RNA expression for biofilm-related targets was determined. (Planktonic: dark gray, pellicle: medium gray, coculture: light gray).

**Figure S3: Biofilm cocultures show increased resistance to colistin compared to planktonic cultures**

S470 WT biofilm cocultures and planktonic cultures were treated with the indicated concentrations of colistin for 2h, and survival was determined by remaining attached fluorescent biomass (cocultures) or live/dead staining (planktonic). Cocultures: solid line. Planktonic cultures: dashed line.

**Figure S4: PaAP mRNA increases over the observed coculture time course**

RNA samples were isolated from S470 WT cocultures at the indicated time points. qRT-PCR was performed to determine the levels of aminopeptidase expression during the time course. RNA levels are shown relative to PaAP expression in planktonic S470 WT cultures during log growth.

**Figure S5: PaAP does not affect pellicle formation under several growth conditions**

Biofilm pellicles were grown in the indicated growth media for 6 hours and biofilm formation was assessed with crystal violet staining. (S470 WT: dark gray, S470 ΔPaAP: light gray). ns, not significant.

**Figure S6: Supernatant fraction aminopeptidase activity**

Aminopeptidase activity of purified cell free supernatants (CFS), vesicle fractions (OMVs), and dense fractions (Heavy) from S470 WT and ΔPaAP OMVs. 10µg total protein was used for each sample.

**Figure S7: WT OMVs can significantly increase Psl/biomass ratios**

Biofilm cocultures were grown, stained for Psl, and quantified. Psl:biomass volume ratios are shown. For treated samples, OMVs were added 1 hpi. Untreated WT and treated ΔPaAP pellicles were stained as above, quantified, and values compared to untreated ΔPaAP pellicles; *p<0.05, **p<0.01, ***p<0.001, ****p<.0001, ns, not significant.

**Figure S8: PaAP^+^ OMVs induce bacterial detachment of pellicles grown in Jensen’s but not in media conditions that do not produce the original PaAP deletion phenotype**

Pellicle biofilms were grown in 96 well dishes for 6 h and biofilms were stained with Crystal Violet. For vesicle treated samples, OMVs were added at 4.5 hpi and imaged at 5 hpi.. Black: S470 WT untreated; dark gray: S470 ΔPaAP untreated; light gray: S470 ΔPaAP + WT OMVs; white: S470 ΔPaAP + ΔPaAP OMVs; *p<0.05, **p<0.01, ***p<0.001, ****p<.0001, ns, not significant.

**Figure S9: Protease inhibitor cocktail does not affect aminopeptidase activity**

S470 WT and ΔPaAP OMVs were treated with a protease inhibitor cocktail (PI) or buffer only and incubated at 37°C. Aminopeptidase activity was measured.

## References

1. Gellatly, S. L. & Hancock, R. E. Pseudomonas aeruginosa: new insights into pathogenesis and host defenses. Pathogens and disease 67, 159–73 (2013).

2. Moradali, M., Ghods, S. & Rehm, B. H. Pseudomonas aeruginosa Lifestyle: A Paradigm for Adaptation, Survival, and Persistence. Frontiers in cellular and infection microbiology 7, 39 (2017).

3. Davey, M. & O’toole, G. Microbial biofilms: from ecology to molecular genetics. Microbiology and molecular biology reviews : MMBR 64, 847–67 (2000).

4. Mann, E. E. & Wozniak, D. J. Pseudomonas biofilm matrix composition and niche biology. Fems Microbiol Rev 36, 893–916 (2012).

5. Laverty, G., Gorman, S. P. & Gilmore, B. F. Biomolecular Mechanisms of Pseudomonas aeruginosa and Escherichia coli Biofilm Formation. Pathogens (Basel, Switzerland) 3, 596–632 (2014).

6. Franklin, M. J., Nivens, D. E., Weadge, J. T. & Howell, L. P. Biosynthesis of the Pseudomonas aeruginosa Extracellular Polysaccharides, Alginate, Pel, and Psl. Front Microbiol 2, 167 (2011).

7. Wang, W., Chanda, W. & Zhong, M. The relationship between biofilm and outer membrane vesicles: a novel therapy overview. FEMS microbiology letters 362, fnv117 (2015).

8. Hoge, R., Pelzer, A., Rosenau, F., Wilhelm, S. & Mendez-Vilas, A. in Current Research, Technology and Education Topics in Applied Microbiology and Microbial Biotechnology number 2, 383–395 (Formatex Research Center).

9. Suleman, L. Extracellular Bacterial Proteases in Chronic Wounds: A Potential Therapeutic Target? Advances in wound care 5, 455–463 (2016).

10. Cahan, R., Axelrad, I., frin, Ohman, D. & Kessler, E. A secreted aminopeptidase of Pseudomonas aeruginosa. Identification, primary structure, and relationship to other aminopeptidases. The Journal of biological chemistry 276, 43645–52 (2001).

11. Galloway, D. Pseudomonas aeruginosa elastase and elastolysis revisited: recent developments. Molecular microbiology 5, 2315–21 (1991).

12. Tielen, P., Rosenau, F., Wilhelm, S., Jaeger, K.-E. E., Flemming, H.-C. C. & Wingender, J. Extracellular enzymes affect biofilm formation of mucoid Pseudomonas aeruginosa. Microbiology (Reading, England) 156, 2239–52 (2010).

13. Bauman, S. J. & Kuehn, M. J. Purification of outer membrane vesicles from Pseudomonas aeruginosa and their activation of an IL-8 response. Microbes and infection 8, 2400–8 (2006).

14. Michel, G. P., Durand, E. & Filloux, A. XphA/XqhA, a novel GspCD subunit for type II secretion in Pseudomonas aeruginosa. Journal of bacteriology 189, 3776–83 (2007).

15. Sarnovsky, R., Rea, J., Makowski, M., Hertle, R., Kelly, C., Antignani, A., Pastrana, D. V. & Fitzgerald, D. J. Proteolytic cleavage of a C-terminal prosequence, leading to autoprocessing at the N Terminus, activates leucine aminopeptidase from Pseudomonas aeruginosa. The Journal of biological chemistry 284, 10243–53 (2009).

16. Orench-Rivera, N. & Kuehn, M. J. Environmentally controlled bacterial vesicle-mediated export. Cellular microbiology 18, 1525–1536 (2016).

17. Kuehn, M. J. & Kesty, N. C. Bacterial outer membrane vesicles and the host–pathogen interaction. Genes & Development 19, 2645–2655 (2005).

18. Cecil, J. D., Sirisaengtaksin, N., O’Brien-Simpson, N. M. & Krachler, A. M. Outer Membrane Vesicle- Host Cell Interactions. Microbiology spectrum 7, (2019).

19. Schooling, S. R. & Beveridge, T. J. Membrane vesicles: an overlooked component of the matrices of biofilms. Journal of bacteriology 188, 5945–57 (2006).

20. Toyofuku, M., Roschitzki, B., Riedel, K. & Eberl, L. Identification of proteins associated with the Pseudomonas aeruginosa biofilm extracellular matrix. Journal of proteome research 11, 4906–15 (2012).

21. Kievit, D. T., Gillis, R., Marx, S., Brown, C. & Iglewski, B. Quorum-sensing genes in Pseudomonas aeruginosa biofilms: their role and expression patterns. Applied and environmental microbiology 67, 1865–73 (2001).

22. Chugani, S. & Greenberg, E. The influence of human respiratory epithelia on Pseudomonas aeruginosa gene expression. Microbial pathogenesis 42, 29–35 (2007).

23. Liberati, N. T., Urbach, J. M., Miyata, S., Lee, D. G., Drenkard, E., Wu, G., Villanueva, J., Wei, T. & Ausubel, F. M. An ordered, nonredundant library of Pseudomonas aeruginosa strain PA14 transposon insertion mutants. P Natl Acad Sci Usa 103, 2833–2838 (2006).

24. Bauman, S. J. & Kuehn, M. J. Pseudomonas aeruginosa vesicles associate with and are internalized by human lung epithelial cells. BMC Microbiol. 9, 26 (2009).

25. Singer, J. T., Phennicie, R. T., Sullivan, M. J., Porter, L. A., Shaffer, V. J. & Kim, C. H. Broad-Host- Range Plasmids for Red Fluorescent Protein Labeling of Gram-Negative Bacteria for Use in the Zebrafish Model System▿ †. Applied and Environmental Microbiology 76, 3467–74 (2010).

26. Moreau-Marquis, S., Redelman, C. V., Stanton, B. A. & Anderson, G. G. Co-culture models of Pseudomonas aeruginosa biofilms grown on live human airway cells. Journal of visualized experiments : JoVE (2010). doi:10.3791/2186

27. Jensen, S., Fecycz, I. & Campbell, J. Nutritional factors controlling exocellular protease production by Pseudomonas aeruginosa. Journal of bacteriology 144, 844–7 (1980).

28. O’Toole, G. A. Microtiter dish biofilm formation assay. Journal of visualized experiments: JoVE (2011). doi:10.3791/2437

29. MacDonald, I. A. & Kuehn, M. J. Stress-Induced Outer Membrane Vesicle Production by Pseudomonas aeruginosa. Journal of Bacteriology 195, 2971–2981 (2013).

30. Ellis, T. N., Leiman, S. A. & Kuehn, M. J. Naturally produced outer membrane vesicles from Pseudomonas aeruginosa elicit a potent innate immune response via combined sensing of both lipopolysaccharide and protein components. Infection and immunity 78, 3822–31 (2010).

31. Finck-Barbançon, V., Goranson, J., Zhu, L., Sawa, T., Wiener-Kronish, J., Fleiszig, S., Wu, C., Mende-Mueller, L. & Frank, D. ExoU expression by Pseudomonas aeruginosa correlates with acute cytotoxicity and epithelial injury. Molecular microbiology 25, 547–57 (1997).

32. Nouwens, A. S., Beatson, S. A., Whitchurch, C. B., Walsh, B. J., Schweizer, H. P., Mattick, J. S. & Cordwell, S. J. Proteome analysis of extracellular proteins regulated by the las and rhl quorum sensing systems in Pseudomonas aeruginosa PAO1. Microbiology (Reading, England) 149, 1311–22 (2003).

33. Zhao, T., Zhang, Y., Wu, H., Wang, D., Chen, Y., Zhu, M. & Ma, L. Z. Extracellular aminopeptidase modulates biofilm development of Pseudomonas aeruginosa by affecting matrix exopolysaccharide and bacterial cell death. Environmental Microbiology Reports 10, 583–593 (2018).

34. Riquelme, S. A., Ahn, D. & Prince, A. Pseudomonas aeruginosa and Klebsiella pneumoniae Adaptation to Innate Immune Clearance Mechanisms in the Lung. Journal of innate immunity 10, 442– 454 (2018).

35. Boles, B. R., Thoendel, M. & Singh, P. K. Rhamnolipids mediate detachment of Pseudomonas aeruginosa from biofilms. Molecular Microbiology 57, 1210–1223 (2005).

36. Petrova, O. E. & Sauer, K. Escaping the biofilm in more than one way: desorption, detachment or dispersion. Current Opinion in Microbiology 30, 67–78 (2016).

37. Zarnowski, R., Sanchez, H., Covelli, A. S., Dominguez, E., Jaromin, A., Bernhardt, J., Mitchell, K. F., Heiss, C., Azadi, P., Mitchell, A. & Andes, D. R. Candida albicans biofilm–induced vesicles confer drug resistance through matrix biogenesis. Plos Biol 16, e2006872 (2018).

38. Hoge, R., Pelzer, A., Rosenau, F. & Wilhelm, S. Weapons of a pathogen: proteases and their role in virulence of Pseudomonas aeruginosa. Current research, technology and education topics in applied microbiology and microbial biotechnology 2, 383–95 (2010).

39. Reichhardt, C., Wong, C., da Silva, D., Wozniak, D. J. & Parsek, M. R. CdrA Interactions within the Pseudomonas aeruginosa Biofilm Matrix Safeguard It from Proteolysis and Promote Cellular Packing. Mbio 9, e01376–18 (2018).

40. Oh, J., Li, X.-H., Kim, S.-K. & Lee, J.-H. Post-secretional activation of Protease IV by quorum sensing in Pseudomonas aeruginosa. Sci Rep-uk 7, 4416 (2017).

41. Jarocki, V. M., Tacchi, J. L. & Djordjevic, S. P. Non-proteolytic functions of microbial proteases increase pathological complexity. Proteomics 15, 1075–88 (2015).

